# Improving Sensitivity of the Digits-in-Noise Test using Antiphasic Stimuli

**DOI:** 10.1101/677609

**Authors:** Karina C. De Sousa, De Wet Swanepoel, David R. Moore, Hermanus Carel Myburgh, Cas Smits

## Abstract

**Objective:** The digits-in-noise test (DIN) has become increasingly popular as a consumer-based method to screen for hearing loss. Current versions of all DINs either test ears monaurally or present identical stimuli binaurally (i.e., diotic noise and speech, N_o_S_o_). Unfortunately, presentation of identical stimuli to each ear inhibits detection of unilateral sensorineural hearing loss (SNHL), and neither diotic nor monaural presentation sensitively detects conductive hearing loss (CHL). Following an earlier finding of enhanced sensitivity in normally hearing listeners, this study tested the hypothesis that interaural antiphasic digit presentation (N_o_S_π_) would improve sensitivity to hearing loss caused by unilateral or asymmetric SNHL, symmetric SNHL, or CHL.

**Design:** This cross-sectional study, recruited adults (18-84 years) with various levels of hearing, based on a four-frequency pure tone average (PTA) at 0.5, 1, 2 and 4kHz. The study sample was comprised of listeners with normal hearing (*n*=41; PTA ≤ 25 dB HL in both ears), symmetric SNHL (*n*=57; PTA > 25 dB HL), unilateral or asymmetric SNHL (*n*=24; PTA > 25 dB HL in the poorer ear) and CHL (*n*=23; PTA > 25 dB HL and PTA air-bone gap ≥ 20 dB HL in the poorer ear). Antiphasic and diotic speech reception thresholds (SRTs) were compared using a repeated-measures design.

**Results:** Antiphasic DIN was significantly more sensitive to all three forms of hearing loss than the diotic DIN. SRT test-retest reliability was high for all tests (ICC *r* > 0.89). Area under the receiver operating characteristics (ROC) curve for detection of hearing loss (> 25 dB HL) was higher for antiphasic DIN (0.94) than for diotic DIN (0.77) presentation. After correcting for age, PTA of listeners with normal hearing or symmetric SNHL was more strongly correlated with antiphasic (*r*_partial_[96]=0.69) than diotic (*r*_partial_=0.54) SRTs. Slope of fitted regression lines predicting SRT from PTA was significantly steeper for antiphasic than diotic DIN. For listeners with normal hearing or CHL, antiphasic SRTs were more strongly correlated with PTA (*r*_partial_[62]=0.92) than diotic SRTs (*r*_partial_[62]=0.64). Slope of regression line with PTA was also significantly steeper for antiphasic than diotic DIN. Severity of asymmetric hearing loss (poorer ear PTA) was unrelated to SRT. No effect of self-reported English competence on either antiphasic or diotic DIN among the mixed first-language participants was observed

**Conclusions:** Antiphasic digit presentation markedly improved the sensitivity of the DIN test to detect SNHL, either symmetric or asymmetric, while keeping test duration to a minimum by testing binaurally. In addition, the antiphasic DIN was able to detect CHL, a shortcoming of previous monaural or binaurally diotic DIN versions. The antiphasic DIN is thus a powerful tool for population-based screening. This enhanced functionality combined with smartphone delivery could make the antiphasic DIN suitable as a primary screen that is accessible to a large global audience.

## INTRODUCTION

Hearing loss presents a significant global health burden as the 4^th^ leading contributor to years lived with disability (Vos et al. 2016). Mounting evidence demonstrates significant associations between hearing loss, depression (Fellinger et al. 2012), unemployment (Ruben 2015), risk for hospitalization (Genther et al. 2013; Reed et al. 2018) and cognitive decline and dementia (Lin et al. 2011; Livingston et al. 2017). Early detection is an essential first step to ameliorate the functional impairment of hearing loss, yet a high proportion of cases remains undetected and untreated (Ki-Moon 2016; Mackenzie and Smith 2009). Contributing to the disparity is lack of routine adult hearing screening programs and rehabilitation options that are either unavailable or prohibitively expensive (Chou et al. 2011; Wilson et al. 2017).

Poor awareness of hearing loss and existing models of clinic-based adult screening among the lay public also contribute to hearing healthcare inaccessibility (Lin et al. 2016). In efforts to increase and decentralize access to detection of hearing loss, screening methods such as the digits-in-noise test (DIN), as an internet or landline phone-based hearing screen have been employed (Smits et al. 2004; Jansen et al. 2010; Watson et al. 2012; Zokoll et al. 2012). The DIN is a speech-in-noise test that uses digit triplets (e.g. 5-9-2), typically presented in steady speech-shaped noise, to measure the speech reception threshold (SRT), expressed in dB signal-to-noise ratio (dB SNR), where a listener can recognize 50% of the digit triplets correctly. Compared to pure tone audiometry or speech recognition in quiet, speech recognition in noise has the advantage of being more characteristic of a person’s hearing ability in real-life situations (Grant et al. 2013). Furthermore, DIN assessment of sensorineural hearing loss (SNHL) correlates highly with pure tone audiometry and eliminates the need for a soundproof booth, calibrated equipment and a test administrator (Smits et al. 2004; Jansen et al. 2010; Potgieter et al. 2016, 2018; Koole et al. 2016).

The DIN was first developed as a national landline telephone test in the Netherlands (Smits et al. 2004) and later also implemented as an internet-based test (Smits et al. 2006). Highly correlated with the audiometric pure tone average (*r*=0.77) it demonstrated sensitivity and specificity of more than 90% to detect sensorineural hearing loss (Smits et al. 2004). Four months after its release, the DIN saw considerable uptake with more than 65,000 tests taken (Smits et al. 2005), demonstrating its role and potential as a large-scale hearing screening tool available to the public. Using simple digits, the test does not require a high degree of linguistic competence (Kaandorp et al. 2016). Various language versions of the DIN have been developed, including British-English (Hall, 2006), American-English (Watson et al. 2012); Polish (Ozimek et al. 2009), French (Jansen et al. 2010) and German (Zokoll et al. 2011).

Despite the success of the DIN in several countries, the need of landline telephones to conduct testing can be problematic, especially in low-and-middle income countries like South Africa where landline penetration is poor (STATSSA 2013). On the other hand, global access to smartphones by adults is estimated to be 80% by the year 2020, providing a modern-day alternative (The Economist 2015). Whereas mobile phone penetration is much higher, the cost to complete the test via a mobile phone call could be more expensive. An alternative is to offer the DIN as a downloadable smartphone application, allowing access to high fidelity broadband signals as opposed to bandwidth signals used in standard telephone networks (Potgieter et al. 2016), and removing the need for cellular connectivity once uploaded. While applicable worldwide, using a mobile platform could potentially address the mostly nonexistent access to hearing screening in low-and-middle income countries. In sub-Saharan Africa, for instance, there is only one audiologist for every million people (Mulwafu et al. 2017). As a result, the South African English DIN was developed and released as the national hearing screening application in 2016, downloadable on iOS and Android smartphones, called hearZA™ (Potgieter et al. 2016; De Sousa et al. 2018). This binaural test version allows for testing under 3 minutes, with high sensitivity (> 80%) to detect SNHL (Potgieter et al. 2016; Potgieter et al. 2018).

There has been a growing interest in increasing the efficiency and sensitivity of existing DINs using various test modifications. Using a fixed-SNR procedure, Smits (2017) showed that the number of digit triplets in a DIN could be reduced to as few as 8 trials, without compromising sensitivity and specificity but sacrificing accurate estimation of the SRT. Furthermore, with the early appearance and high prevalence of high frequency hearing loss, use of low-pass filtered masking noise to improve sensitivity of the DIN to high frequency hearing loss has been investigated, showing either higher (Vlaming et al. 2014) or similar (Vercammen et al. 2018) area under the receiver operating characteristics (ROC) curve compared to DINs with standard speech-shaped noise. Therefore, when using homogenized digits to ensure high test-retest reliability, these modifications could make the DIN test more applicable to persons with noise-induced or age-related hearing loss (Vlaming et al. 2014).

Current versions of all DINs either sequentially test each ear (monaurally) or present the test stimuli binaurally and identically to each ear (homophasic or diotic). This binaural DIN setup allows for rapid testing in approximately 3 minutes, whereas sequential testing of each ear doubles test time and may thus reduce uptake and completion. Using diotic presentation may, however, preclude detection of unilateral or asymmetric sensorineural hearing loss (SNHL). These listeners may pass the diotic DIN test because performance is largely based on the functionally better ear (Potgieter et al. 2018). Furthermore, both monaural and diotic testing is insensitive to the attenuation caused by conductive hearing loss (CHL) because most DINs are presented at suprathreshold intensities. To improve the sensitivity of the DIN, especially for listeners with unilateral, asymmetrical SNHL and CHL, this study evaluated the use of a DIN test paradigm using digits that are phase inverted (antiphasic) between the ears, while leaving the masking noise interaurally in-phase. Such a configuration of stimuli (N_o_S_π_) was shown to improve DIN SRTs in normal hearing listeners (Smits et al. 2016).

Sensitivity differences between diotic and antiphasic auditory stimulus presentations are commonly known as the binaural masking level difference (Hirsh 1948). Before the widespread use of the auditory brainstem response, binaural masking level difference was employed to distinguish between different types of hearing loss (Olsen et al. 1976; Wilson et al. 2003). Binaural masking level difference was reported to be poorer for listeners with various types and configurations of hearing loss compared to normal hearing controls. Wilson and colleagues (1985) investigated speech masking level difference for people with unilateral SNHL. In the diotic condition (N_o_S_o_), only slight SNR variations were observed across a range of interaural level differences. However, in the antiphasic condition (N_o_S_π_), SNRs became worse with increasing interaural level differences.

Smits and colleagues (2016) examined SRTs in different listening conditions for the Dutch and American English DIN among normal hearing listeners. Results indicated that the threshold advantage over monotic presentation provided by diotic (N_o_S_o_) presentation was small (≅ 1 dB). However, the use of antiphasic digits (N_o_S_π_) provided a further ≅ 5 dB advantage. Listeners with unilateral SNHL or CHL are not expected to have full access to the antiphasic advantage due to subtle timing irregularities caused by peripheral hearing loss, either sensorineural (Jerger et al. 1984; Thornton et al. 2012; Wilson et al. 1985) or conductive (Hartley and Moore 2003; Jerger et al. 1984). In cases of symmetric hearing loss, the antiphasic advantage is expected to decrease as the degree of hearing loss increases because of increasing threshold and timing cue deterioration (Wilson et al. 1994). These findings support the idea that antiphasic digit presentation could sensitize the DIN for a wider range of hearing loss types while using a single binaural test. This would improve the function of current consumer-based DINs.

The objective of this study was, therefore, to determine whether antiphasic digit presentation improves detection of hearing loss relative to the diotic presentation.

## MATERIALS AND METHODS

### Study design and participants

A cross-sectional, repeated-measure study of the DIN SRT comparing diotic and antiphasic presentation within and between listeners of varying types and degrees of hearing loss was conducted. Listeners were recruited from a student population, a University clinic, and hospital and private practices in the Gauteng province of South Africa. Adults (18-84 years; Table 1) with various levels of hearing were recruited, based on a four frequency (0.5,1,2 and 4kHz) pure tone average (PTA). The study sample included normal hearing (*n*=41; pure tone average (PTA) ≤ 25 dB HL in both ears), symmetric SNHL (*n*=57; PTA > 25 dB HL) and unilateral or asymmetric SNHL (*n*=24; PTA > 25 dB HL in the poorer ear). The better ear PTA of listeners with asymmetric SNHL did not exceed 45 dB HL. A sample of listeners with CHL (*n*=23; PTA > 25 dB HL and PTA air-bone gap ≥ 20 dB HL in the poorer ear) was also recruited, including 3 listeners with symmetric and 20 with unilateral or asymmetric hearing loss. Bone conduction pure tone average thresholds (0.5,1,2,4 kHz) for the poorer ear did not exceed 25 dB HL, except for one listener with CHL with poorer ear bone conduction PTA of 28 dB HL. Asymmetric hearing loss was defined as an interaural difference >10 dB (PTA). Hearing sensitivity categories were based on poorer ear PTA and categorized as excellent (0-15 dB HL), minimal (16-25 dB HL), mild (26-40 dB HL), moderate (41-55 dB HL) and severe-to-profound (56-120 dB HL). For analyses, the ‘excellent’ and ‘minimal’ categories were combined into a single ‘normal’ category. Listeners had various levels of English-speaking competence. Non-native English speakers self-reported their level of competence on a non-standardized scale from 1-10, a higher score indicating better competence (Potgieter et al. 2018).

**Table 1.** Analysis of variance statistics for the effect of test presentation and category of hearing loss.

|  |  |  |  |  |
| --- | --- | --- | --- | --- |
|  | **df** | **F** | **Sig** | **Partial Eta Squared** |
| Test Type<br>(Diotic vs Antiphasic) | 1, 141 | 497.06 | < .001 | 0.78 |
| Hearing Category | 3,141 | 31.88 | < .001 | 0.41 |
| Test Type*Hearing Category | 3, 141 | 57.81 | < .001 | 0.55 |

The Health Sciences Research Ethics Committee, University of Pretoria approved the study protocol (number 58/2017). All eligible participants were informed on the study aims and procedures and provided consent before participation.

### Procedures and Equipment

The smartphone application for the South African English DIN was adapted for antiphasic stimulus presentation. Original homogenized diotic digits were phase reversed for antiphasic presentation. The phase inversion was completed in MatLab by multiplying each sample in one channel of the digit triplet sound file by −1. The DIN application was designed in Android Studio version 2.3.0 and written in Java version 1.8.0, consistent with the original *hearZA* App. The application stored a list of 120 different digit-triplets, randomly selected for presentation at the beginning of each test (Potgieter et al. 2016). Randomized triplet selection was done with replacement, meaning that the same triplet could occur more than once in one test. Triplets were presented with 500 ms silent intervals at the beginning and end of each digit-triplet. Successive digits were separated by 200 ms of silence with 100 ms of jitter (Potgieter et al. 2016). The test used a fixed noise level and variable speech level when triplets with negative SNRs were presented. To prevent clipping of the signal, the speech level was fixed, and the noise level varied once the SNR became positive (Potgieter et al. 2016). The speech-weighted masking noise was delivered interaurally in-phase, and the digits were either in-phase (diotic; N_o_S_o_) or were phase inverted between the two ears (antiphasic; N_o_S_π_). To prevent possible learning of the masking noise (Lyzenga and Smits 2011), noise ‘freshness’ was ensured for each trial by creating a long noise file and selecting successive fragments from a random offset within the first 5 seconds. Both diotic and antiphasic versions of the DIN consisted of 23 digit-triplets. The SNR varied in fixed step sizes (4 dB SNR for the first three steps, thereafter continuing in 2 dB steps) starting at 0 dB SNR using a one-up one-down staircase procedure, tracking the SNR at which 50% of the digit triplets were correctly identified (Smits et al. 2004; Potgieter et al. 2016). For the first three steps, SNR became progressively more negative by 4 dB per step for correct responses but increased by 2 dB per step for incorrect responses. A digit-triplet was only considered correct when all digits were entered correctly. The SRT was calculated by averaging the last 19 SNRs, in line with the currently used *hearZA* test.

After completion of pure tone audiometry, participants completed five DIN tests, each lasting about 3 minutes, on a Samsung Trend Neo smartphone coupled with manufacturer supplied (wired) earbuds in a quiet, office-like room. The first training test used antiphasic presentation. The remaining four DIN tests alternated between antiphasic and diotic DIN, with a test and retest for each participant. The test order was therefore: (1) antiphasic training list, (2) antiphasic test, (3) diotic test, (4) antiphasic re-test and (5) diotic re-test.

### Statistical Analysis

Statistical analysis was done using the Statistical Package for the Social Sciences (IBM SPSS v25.0). A sample size of 122 listeners (24 with normal hearing PTA ≤ 25 dB HL, 24 with asymmetric hearing loss and 74 with either symmetric normal hearing PTA ≤ 25 dB HL or symmetric sensorineural hearing loss with PTA ≥ 26 dB HL) would provide a medium effect size (Cohen’s *f* = 0.25), with 80% statistical power at two-tailed significance level of 0.05, to test both hypotheses. The sample of 23 listeners with CHL was subsequently added.

The effect of test condition (i.e., diotic or antiphasic) and hearing loss category (i.e., type and symmetry of hearing loss) on the SRT was assessed using repeated-measures analysis of variance. Post hoc comparisons used Bonferroni adjustment for multiple comparisons. In cases where sphericity was violated, Greenhouse-Geisser corrections were applied. Analysis of covariance was used to determine effects of age and English-speaking competence on the diotic and antiphasic SRT. General linear regression was used to test whether the slope of the relation between PTA and SRT differed between antiphasic and diotic testing. The effect of test repetition on antiphasic SRT was investigated using a paired sample *t*-test. Intraclass correlation coefficients (ICC) were calculated and were based on a mean rating of the number of observations (i.e. test and retest; *k*=2) of both diotic and antiphasic test conditions, absolute agreement, and a two-way mixed-effects model. In addition, measurement error between test-retest for diotic and antiphasic presentation was calculated by determining quadratic mean 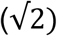 of within-subject standard deviations for the test-retest measures. All subsequent analyses were conducted by averaging the test and retest SRT values for the diotic and antiphasic DIN. Associations between poorer ear PTA and SRT were examined using Pearson’s partial correlation. Receiver operating characteristics (ROC) curves were calculated to determine the sensitivity and specificity of the DIN tests for different cutoff values, to detect mild hearing loss and worse (PTA > 25 dB HL) and moderate hearing loss and worse (PTA > 40 dB HL). SRT cut-off values corresponding to reasonably high sensitivity and specificity were chosen, while demonstrating the trade-off between sensitivity and specificity (i.e. higher sensitivity with consequent lower specificity).

## RESULTS

Listeners with normal hearing had lower SRTs than those with hearing loss using both diotic and antiphasic testing (Fig. 1). However, antiphasic testing was significantly more sensitive to all three forms of hearing loss than diotic testing (Table 1).

**Figure 1.**
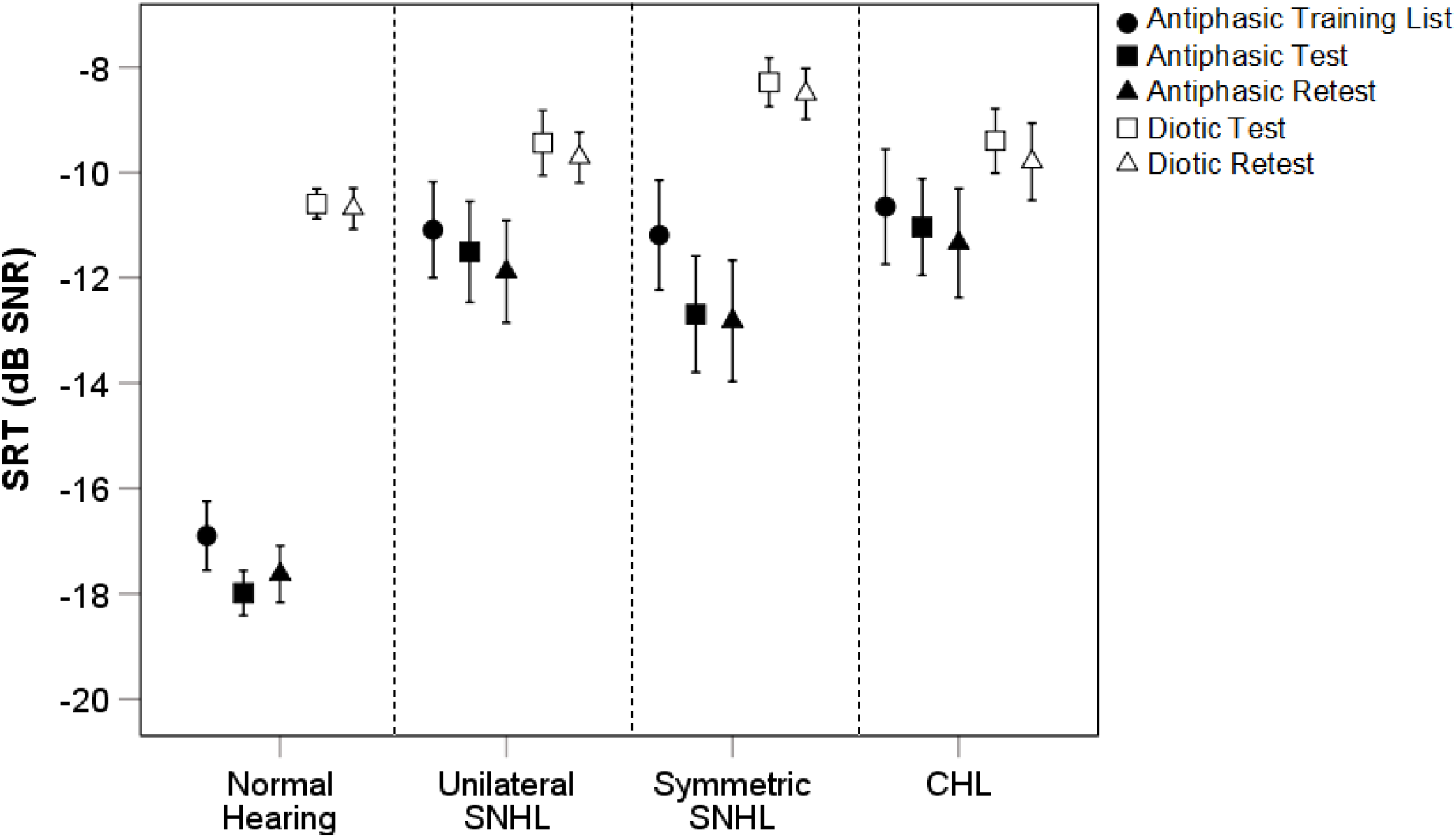
Antiphasic and diotic SRT according to hearing category. SRT; speech reception threshold, dB; decibel, HL; hearing level. Error bars are 95% confidence intervals.

Across all hearing categories, after controlling for age, poorer ear PTA was significantly correlated to both diotic and antiphasic SRT (*p* < 0.001). The correlation was, however, stronger for antiphasic (*r*_partial_[145]=0.82] than diotic SRT (*r*_*partial*_[145]=0.44). For listeners with either normal hearing or symmetric SNHL, poorer ear PTA was significantly (*p* < 0.001) correlated with both antiphasic (*r*_*partial*_[96]=0.69) and diotic (*r*_*partial*_[96]=0.54) SRTs (Fig. 2). However, the slope of the fitted regression was significantly steeper for antiphasic SRTs (*t*(1)=7.79.14, *p* < 0.001). Antiphasic SRTs of listeners with normal hearing or CHL, were more strongly correlated to poorer ear PTA (*r*_*partial*_[62]=0.92) than diotic SRTs (*r*_*partial*_[62]=0.54). The slope of the fitted regression was also significantly steeper for antiphasic compared to diotic SRTs (*t*[1]=11.84, *p* < 0.0001), indicative of greater sensitivity of the antiphasic DIN. The severity of unilateral or asymmetric SNHL (poorer ear PTA) was unrelated to SRT. For the diotic DIN, there was substantial overlap between the SRTs of normal hearing listeners and those in each of the three hearing loss groups (Fig. 2A), even for PTAs in the moderate or greater hearing loss ranges (Table 2). The SRT overlap was less substantial for the antiphasic DIN, with listeners with mild poorer ear hearing loss corresponding in SRTs to the normal hearing group (Fig. 2B).

**Figure 2.**
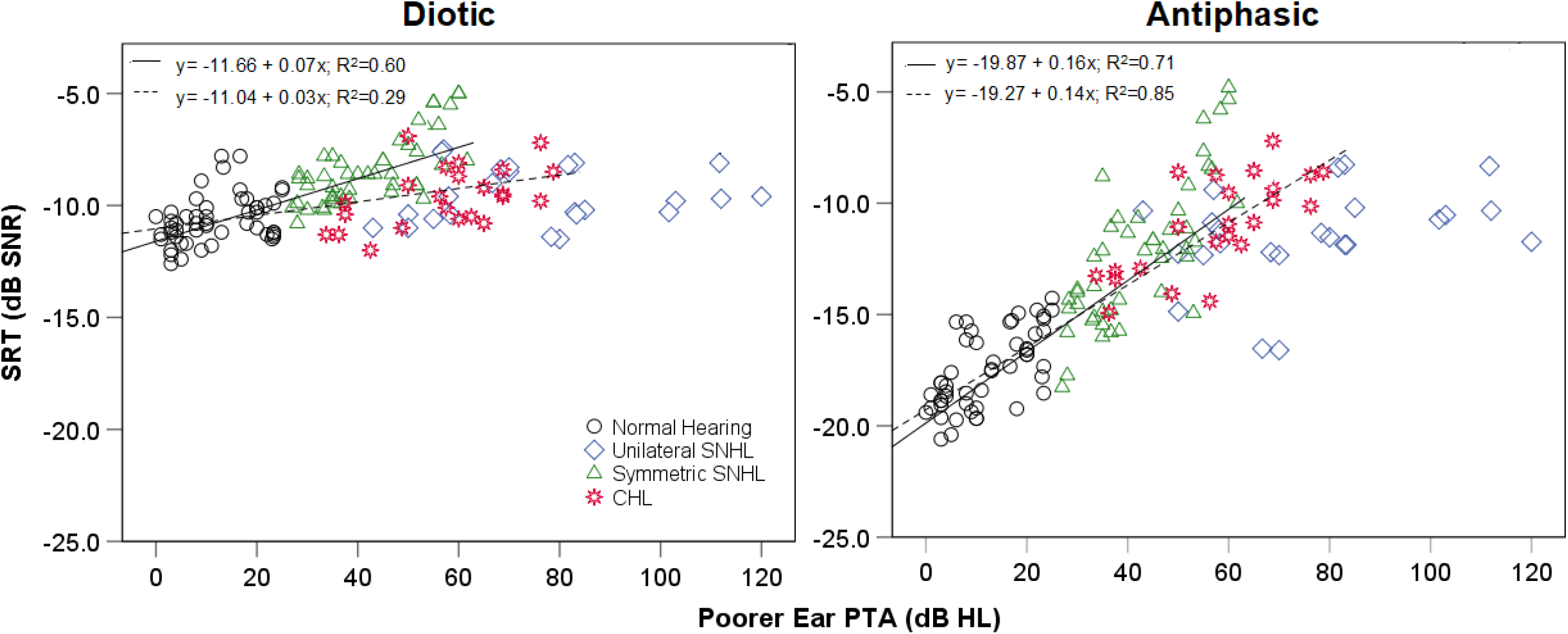
Correlations of the diotic and antiphasic DIN to poorer ear PTA. Solid lines are regression lines fitted to normal hearing and symmetric SNHL group data. Dashed lines are regression lines fitted to normal hearing and CHL group data. SRT; speech reception threshold, dB; decibel, SNR; signal to noise ratio, PTA; pure tone average, HL; hearing level.

**Table 2.** Diotic and antiphasic DIN SRT for listeners with normal hearing, symmetric SNHL, unilateral or asymmetric SNHL and CHL according to PTA hearing loss categories.

|  |  |  | Excellent<br>(0-15 dB) | Minimal<br>(16-25 dB) | Mild<br>(26-40 dB) | Moderate<br>(41-55 dB) | Severe-<br>profound<br>(56-120 dB) |
| --- | --- | --- | --- | --- | --- | --- | --- |
| **NH & SHL** |  | *n* | 26 | 15 | 23 | 24 | 10 |
|  |  | Age Range (Years) | 19-67 | 23-84 | 39-84 | 51-84 | 67-79 |
|  | Diotic DIN | Mean SRT(SD) | -11.1 (0.8) | -9.7 (1.1) | -10 (1.1) | -8.7 (0.9) | -6.4 (1.5) |
|  |  | SE | 0.16 | 0.28 | 0.22 | 0.19 | 0.49 |
|  | Antiphasic DIN | Mean SRT(SD) | -18.4 (1.4) | -16.7 (1.6) | -15.7 (1.8) | -12.4 (2.1) | -8.2 (2.7) |
|  |  | SE | 0.28 | 0.41 | 0.37 | 0.43 | 0.85 |
| **UHL** |  | *n* | 0 | 0 | 0 | 4 | 20 |
|  |  | Age Range (Years) | - | - | - | 25-63 | 18-72 |
|  | Diotic DIN | Mean SRT(SD) | - | - | - | -10.8 (0.3) | -9.4 (1.2) |
|  |  | SE | - | - | - | 0.15 | 0.27 |
|  | Antiphasic DIN | Mean SRT(SD) | - | - | - | -12.5 (1.9) | -11.3 (2.2) |
|  |  | SE | - | - | - | 0.93 | 0.49 |
| **CHL** |  | *n* | 0 | 0 | 4 | 4 | 15 |
|  |  | Age Range (Years) |  |  | 18-44 | 19-62 | 20-68 |
|  | Diotic DIN | Mean SRT(SD) | - | - | -10.7 (0.7) | -9.8 (2.3) | -9.3 (1) |
|  |  | SE | - | - | 0.35 | 1.12 | 0.27 |
|  | Antiphasic DIN | Mean SRT(SD) | - | - | -13.7 (0.9) | -11.7 (2.4) | -10.1 (1.8) |
|  |  | SE | - | - | 0.43 | 1.19 | 0.46 |

*DIN: digits-in-noise, NH; Normal Hearing, SHL; Symmetric SNHL, UHL; unilateral sensorineural hearing loss, CHL; conductive hearing loss*

ROC analysis, including poorer ears of all participants, (Fig. 3) showed higher areas under the curve for antiphasic DIN compared to diotic DIN to detect PTA >25 dB HL (0.95; 95% CI, 0.91 to 0.98 vs 0.78; 95% CI, 0.69 to 0.86) and >40 dB HL (0.96; 95% CI, 0.93 to 0.99 vs 0.80; 95% CI, 0.73 to 0.87). Antiphasic DIN was, therefore, more sensitive and specific to hearing loss (of either type and symmetry) compared to diotic DIN. SRT cut-offs in Table 3 demonstrate the trade-off between sensitivity and specificity to detect PTA > 25 dB HL and > 40 dB HL.

**Figure 3.**
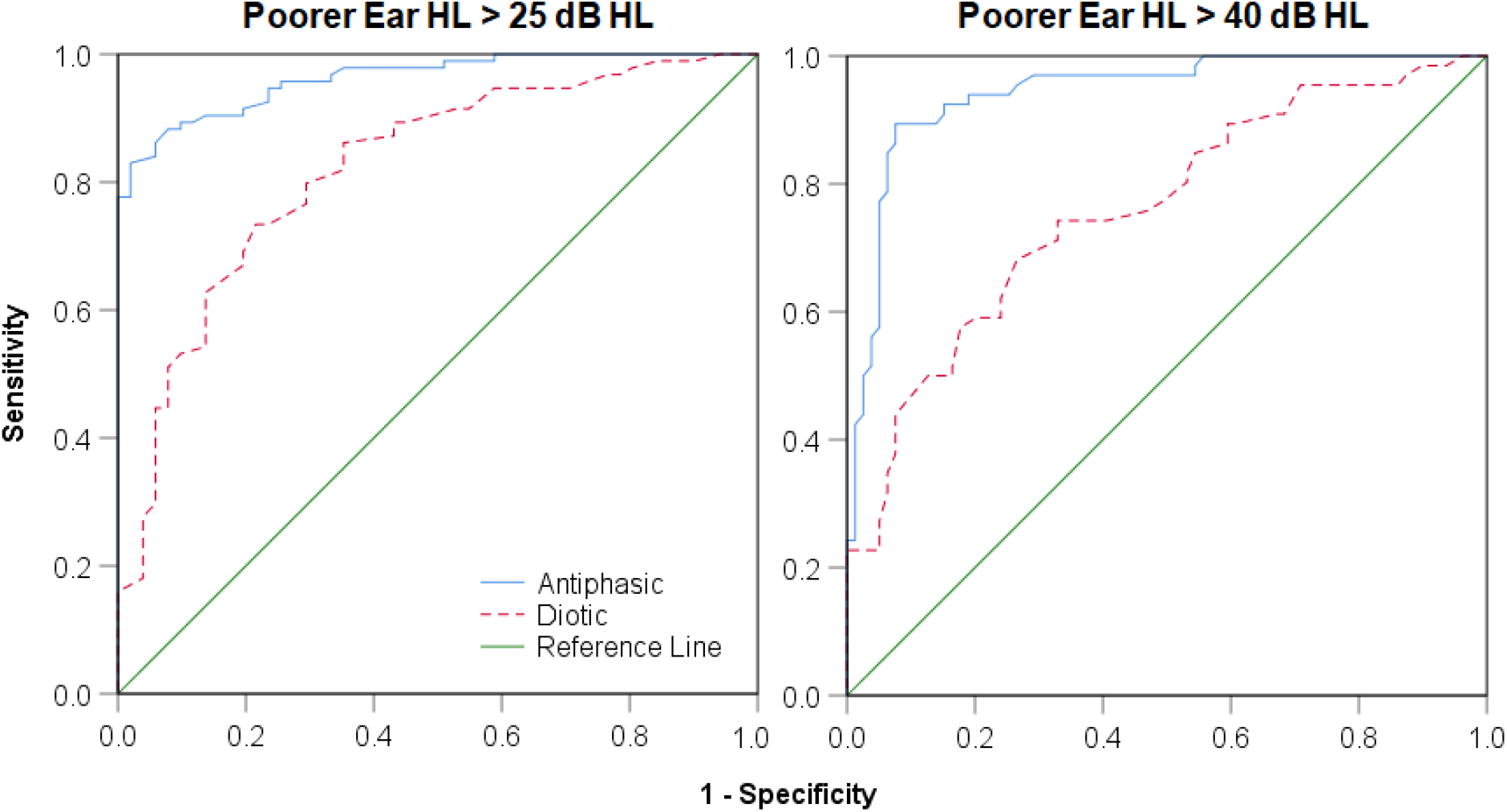
ROC curves presenting test characteristics of the antiphasic and diotic DIN for detecting poorer ear HL > 25 dB HL (left) and > 40 dB HL (right) in all participants with normal hearing, unilateral SNHL, symmetric SNHL and CHL.

Antiphasic DIN test repetition produced a significant mean SRT improvement (0.9 dB SNR; 95% CI 0.61 to 1.4) across hearing categories following the presentation of the initial antiphasic training list (*t*[144]=5.1, *p* < 0.001; Fig. 1). However, between the subsequent test and retest, the mean SRT difference was not significant (*p =* 0.86).

Similarly, diotic test and retest showed no significant SRT difference (*p* = 0.6). SRT test-retest reliability was high for listeners with normal hearing or SNHL for both diotic DIN (ICC=0.89; 95% CI 0.85 to 0.93) and antiphasic DIN (ICC=0.94; 95% CI, 0.91 to 0.96). Listeners with CHL had high test-retest reliability for antiphasic DIN, with ICC of 0.88 (95% CI, 0.72 to 0.95; *p* < 0.001), but had poorer ICC of 0.61 for homophasic DIN (95% CI, 0.09 to 0.83; *p* < 0.05). Diotic DIN had lower measurement error (1.1 dB; 95% CI 0.9 to 1.2) than the antiphasic DIN (1.4 dB; 95% CI 1.2 to 1.5) for the whole sample, but the variance between listeners was much higher for the antiphasic DIN than for the diotic DIN (Table 2).

The effect of competence in the English language on SRT was assessed by dividing listeners into high competence (>7; *n*=73) and lower competence (≤7; *n*=72) groups. Controlling for poorer ear PTA and age, no significant SRT difference (*p* = 0.16) was found between the two groups for either the diotic DIN (*F*[1,141]=2.47, partial η^2^=0.02) or the antiphasic DIN (*F*[1,141]=1.98, partial η^2^ = 0.02).

## DISCUSSION

Antiphasic presentation improved the test characteristics of the smartphone DIN test with higher sensitivity and specificity to detect hearing loss of various degrees, types and symmetries than the diotic DIN. With monaural testing, it is possible to segregate a ‘better’ ear from a ‘poorer’ ear. Traditionally, emphasis has been placed on the function of the ‘better’ ear to assess activity and participation, but there is now considerable evidence that asymmetric or unilateral hearing loss can reduce these aspects of hearing health almost, or as much as symmetric binaural HL (Firszt et al. 2015; Rothpletz et al. 2012; Vannson et al. 2015). It is thus important to assess the function of both ears, working together. Binaural tests, as used here, are more dependent on the relative function of both ears (Supplementary Figure) but, because of interaural summation and unmasking effects that interaction is complex (Hall et al., 1995, 1998). A screening test should be rapid, is not intended to be diagnostic, and persons who fail the test must be referred for diagnostic testing (Wilson and Jungner, 1968). The antiphasic DIN is a rapid test compared to sequential monaural testing and aims to detect all hearing losses that require further diagnostic assessment.

### Mechanisms of antiphasic advantage

Listeners with normal hearing in both ears were at a significant advantage for understanding speech-in-noise compared to listeners with either type or symmetry of hearing loss. This advantage is due to several mechanisms, but the primary one is binaural integration. In spatial hearing, when sound from a lateral source arrives at the nearer ear earlier than the far ear, interaural phase differences are processed as spatial cues. Brainstem neurons detect interaural timing differences as small as 10 μs (Brughera et al. 2013), equal to about 2° of space (Middlebrooks and Green 1991). In the antiphasic DIN, the 180° interaural phase difference of the digits simulates an interaural timing difference, separating virtually the target speech from the noise. We introduced a phase inversion in the speech signals between the ears, leaving the noise in-phase (N_o_S_π_) since the SRT improvement is larger compared to the N_π_S_o_ condition (Olsen et al. 1976). Listeners in our study with ‘normal’ hearing had 6-8 dB better antiphasic than diotic SRT, in line with the study of Smits et al. (2016). Peripheral hearing loss disrupts interaural timing differences by desynchronizing neural activity from the affected ear(s), reducing the antiphasic advantage (Jerger et al. 1984; Welsh et al. 2004; Vannson et al. 2017). Predicted poorer diotic SRTs due to loss of outer hair cell function and associated cochlear compression were also observed for listeners with symmetric SNHL. Antiphasic SRTs, however, demonstrated greater threshold differences in listeners with symmetric SNHL between the various categories of hearing sensitivity compared to diotic SRT.

### Unilateral hearing loss

Diotic presentation in unilateral SNHL does not result in strongly elevated SRTs compared to listeners with bilateral normal hearing, because performance mainly reflects the better ear. Furthermore, the 1 dB advantage provided by binaural summation (Smits et al. 2016), was ineffective to detect unilateral SNHL. Listeners in this study, with moderate unilateral SNHL achieved diotic SRTs comparable to listeners with normal hearing. Similarly, diotic SRTs of those with severe-to-profound unilateral or asymmetric SNHL compared to those with only mild symmetric SNHL. Since listeners with unilateral SNHL could only adequately hear the digits presented to the better ear, binaural interaction was either minimal or entirely absent. Antiphasic SRTs were, as expected, significantly poorer and better reflected the degree of hearing loss in the poorer ear than did diotic SRTs.

Listeners with strongly asymmetric hearing loss could increase the overall presentation level of the DIN test by self-selecting a higher listening level. Some of these listeners may then have enough residual hearing in the poorer ear to achieve a degree of binaural advantage in antiphasic conditions when the signal intensity is brought to threshold in that ear. However, the degree to which overall level adjustment compensates for asymmetric hearing loss is also restricted to the tolerance of masking noise in the better ear (Jerger et al. 1984). Three listeners with primarily high-frequency unilateral SNHL had antiphasic SRTs within the normal range. Since interaural timing differences are low frequency (< 1500 Hz) dependent (Middlebrooks and Green 1991), it is expected that the favourable antiphasic SRTs obtained in these three listeners was due to involvement of their residual low-frequency hearing.

### Conductive hearing loss

The antiphasic test paradigm was very successful in detecting listeners with CHL. A person with symmetric CHL could overcome loudness attenuation of the standard diotic signals by increasing the overall presentation level, thereby achieving near-normal standard SRTs, as seen in listeners with mild and moderate CHL. Diotic SRTs were slightly poorer across consecutive hearing sensitivity categories (mild, moderate and severe-to-profound), but in most cases (20/23) were still within the normal hearing range. Earlier studies demonstrated that antiphasic processing is disrupted by acute CHL that both attenuates and delays sound passing through the ear (Hartley and Moore 2003). Chronic CHL commencing in infancy can impair antiphasic listening even after CHL has resolved (Moore et al. 1991; Pillsbury et al. 1991) and produces a number of neurological changes affecting binaural integration (Polley et al. 2013). Due to the disruption in interaural timing difference caused by CHL, antiphasic SRTs in our study deviated considerably from listeners with normal hearing, in contrast to diotic SRTs.

### Training and reliability

Listeners with normal hearing and SNHL had a small training effect between the antiphasic training list and test condition. There were no significant SRT differences between the diotic DIN and antiphasic DIN test and retest measurements. Similar findings were reported by Smits et al. (2013), suggesting that SRT improvement from the training list to the first test condition is due to a procedural learning effect in naïve listeners. Overall, the antiphasic DIN test-retest reliability was high and better detected CHL as opposed to diotic SRTs. Overall, across the entire sample in this study, antiphasic DIN test characteristics for detecting mild and moderate hearing loss was high. The area under the receiver operating characteristics curve for antiphasic test accuracy for hearing losses of >25 dB HL and >40 dB HL was significantly higher (0.94 and 0.96) than for diotic testing (0.78 and 0.80).

### Clinical implications

The high sensitivity of a 3-minute antiphasic DIN to detect hearing loss of various types, symmetries and degrees holds significant potential for population-based screening. CHL, in the form of otitis media, is typically more prevalent among underserved, remote and poor populations than other forms of hearing loss (Hunter et al. 2007; Cameron et al. 2014) but is not easily detected with currently used DIN tests. Since the DIN can be used in children as young as 4 years of age (Koopmans et al. 2018), the antiphasic DIN test may be a means of early identification in those populations, once age-specific normative SRT scores are established. School-aged screening programs where the DIN has already been successfully implemented (Denys et al. 2018), could similarly benefit from an antiphasic variant to improve sensitivity and reduce test duration from a monaural to a binaural test. Of course, the completion of a single antiphasic DIN test would not be able to differentiate between either CHL or SNHL, or as with monaural testing, between unilateral or bilateral hearing loss. However, following up on initial screening with other DIN variants (e.g. monaural, filtered or modulated noise) for those who fail the antiphasic test could potentially allow for categorization into bilateral, unilateral or CHL.

A smartphone platform of test delivery has proved a successful method of screening, allowing for directed referrals from cloud-based data management platforms (De Sousa et al. 2018), thereby optimizing resource allocation. Furthermore, it has been shown that the test can be done reliably across various smartphone devices (either iOS or Android operated) and transducers (Potgieter et al., 2016; De Sousa et al. 2018). Analysis of the *hearZA* tests taken approximately a year and a half after its release, showed high test uptake (> 30 000 tests), especially among an important target population of users younger than 40 years (De Sousa et al. 2018). The development of the antiphasic DIN test in other language variants, however, is recommended to make it accessible to a large global audience.

In conclusion, antiphasic SRTs correlated significantly better to poorer ear PTA than diotic SRTs. As a result, antiphasic presentation markedly improved sensitivity to detect SNHL and CHL, either symmetric or asymmetric, making it a powerful tool for population-based screening.

## ACKNOWLEDGEMENTS

The authors thank all the participants of this study, Steve Biko Academic Hospital and all participating private practices for their assistance with data collection. The authors thank Li Lin for assistance with data analysis.

This research was funded by the National Institute of Deafness and Communication Disorders of the National Institutes of Health under award number 5R21DC016241-02. Additional funding support was obtained from the National Research Foundation (Grant PR_CSRP190208414782). The 2^nd^ author’s relationship with the hearX Group and hearZA includes equity, consulting, and potential royalties. The last author’s relationship with the hearX Group and hearZA includes equity, consulting and potential royalties. David Moore is supported by Cincinnati Children’s Research Foundation and by the NIHR Manchester Biomedical Research Centre.

