## Supplemental Figure 1 for "Improving Sensitivity of the Digits-in-Noise Test using Antiphasic Stimuli"

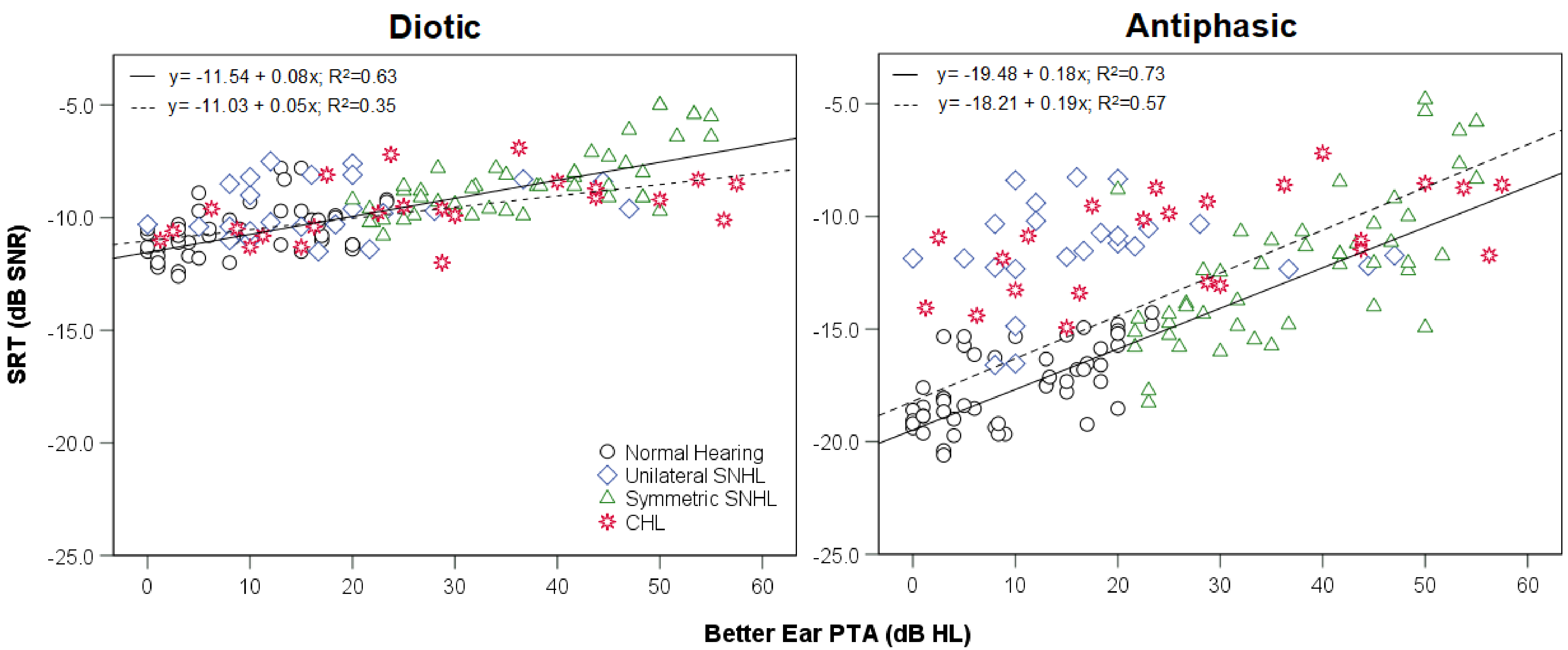


**Figure. Correlations of the diotic and antiphasic DIN to better ear PTA.** Solid lines are regression lines fitted to normal hearing and symmetric SNHL group data. Dashed lines are regression lines fitted to normal hearing and CHL group data. SRT; speech reception threshold, dB; decibel, SNR; signal to noise ratio, HL; hearing level.
